# *In silico* analysis of coding SNPs and 3′-UTR associated miRNAs in *DCAF17* gene that may affect the regulation and pathogenesis of Woodhouse-Sakati Syndrome

**DOI:** 10.1101/601310

**Authors:** Abdelrahman H. Abdelmoneim, Asia M. Elrashied, Alaa I. Mohammed, Sara A. Mirghani, Rania E. Osman, Esraa O. Gadim, Mohamed A. Hassan

## Abstract

**Background:** Woodhouse-Sakati Syndrome refers to a group of inherited disorders characterized by alopecia, hypogonadism, diabetes mellitus, hypothyroidism and progressive extrapyramidal signs. The aim of this study is to identify the pathogenic SNPs in the *DCAF17* gene with their related mciroRNAs and their effect on the structure and function of the protein.

**Material and Methods:** We used different bioinformatics tools to predict the effect of each SNP on the structure and function of the protein. After that we defined the miRNAs founded in the 3′-UTR region on the *DCAF17* gene and studied the annotations relative to it.

**Results:** Ten deleterious SNPs out of 339 were found to have a damaging effect on the protein structure and function, with one significant micoRNA in the 3′-UTR region.

**Conclusion:** This was the first in silico analysis of *DCAF17* gene, in which 10 novel mutations were found using different bioinformatics tools that could be used as a diagnostic markers for Woodhouse-Sakati syndrome, with one relevant microRNA that can regulate the function of the protein.

## 1. Introduction

Woodhouse-Sakati Syndrome (WSS, MIM: 241080) is a rare autosomal-recessive multi systemic disorder which is characterized by a combination of hypogonadism, alopecia, Diabetes Mellitus (DM), mental retardation and extrapyramidal signs.^(1–10)^

It was originally described in a number of consanguineous Saudi families in the Middle East, but has recently been reported in other ethnicities as well.^(11, 12)^ Since its original description in 1983, approximately 50 cases have been reported until now.^(13)^ Exogenous hormone therapy promoting secondary sex characteristic development and insulin are among the recommended treatments for the disease.^(14, 15)^

The availability of vast amounts of sequence data, coupled to advances in computational biology in the recent years provides an ideal framework for in silico gene expression analysis. Single Nucleotide Polymorphisms (SNPs) make up about 90% of DNA sequence variations, making it the most common type of genetic variation. The SNPs that are most likely to have a direct impact on the protein product of a gene are those that change the amino acid sequence and variants in gene regulatory regions, which control protein expression levels.(16-18)

The disease was first described in 2008 by al Alazami et al,^(4, 11)^ Mutations in the *DCAF17* gene are the cause of Woodhouse-Sakati syndrome.^(3, 19, 20)^ It is located on chromosome 2q31.1.^(4, 21)^ it encodes a nucleolar protein with poorly understood function, adding to that the pathogenic mechanism underlying WSS is also un known.^(13)^

The aim of this study was to identify the pathogenic SNPs in *DCAF17* gene located in the coding region and analyzing the microRNA(miRNA) in the 3′-untranslated regions (3′-UTR) using in silico prediction software’s that determine the structure, function and regulation of their respective proteins. This is the first *in silico* analysis of it is kind in *DCAF17* gene to address this matter.

## 2. Material and Methods

### 2.1. Retrieving nsSNPs

SNPs associated with *DCAF17* gene were obtained from the Single Nucleotide Polymorphism database (dbSNP) in the National Center for Biotechnology Information (NCBI). (http://www.ncbi.nlm.nih.gov/snp/).

The sequence and natural variants of DCAF17 protein were obtained from the UniProt database as it considered as the most reliable and unambiguous database for protein sequences.^(22)^ (https://www.uniprot.org/).

A total number of 339 nonsynonymous Single Nucleotide Polymorphisms (ns SNPs) were found from NCBI database, all of them were subjected to in silico analysis using ten different algorithms and softwares; SIFT, PROVEAN, PolyPhen-2, SNPs&GO, PhD-SNP, PMUT, I-mutant, Project Hope, GeneMANIA and Chimera.

### 2.2. Identifying the most damaging nsSNPs and disease related mutations

Five different bioinformatics tools were utilized to predict functional effects of nsSNPs obtained from dbSNP database on the protein. These algorithmic include: SIFT, PROVEAN, PolyPhen-2, SNAP2 and PhD-SNP.

The SNPs predicted deleterious by at least four softwares were considered high risk nsSNPs and investigated further.

#### 2.2.1. SIFT Server

Phenotypic effects of amino acid substitution on protein function were predicted by using Sorting Intolerant From Tolerant (SIFT) server, which is a powerful tool used to fulfill this purpose. A list of nsSNPs (rsIDs) from NCBI’s dbSNP database was submitted as a query (original) sequence to SIFT to predict tolerated and deleterious substitutions for every position of a protein sequence. The server divides the results into “Deleterious” and “Tolerated”, nsSNPs with SIFT score ≤ 0.05 were classified as deleterious and are further analyzed to identify the damaging ones, and those > 0.05 were classified as tolerated and are not further analyzed. (Available at: http://sift.bii.a-star.edu.sg/).^(23–26)^

#### 2.2.2. Provean Server

(Protein Variation Effect Analyzer) is the second software tool used. It also predicts the effect of an amino acid substitution on the biological function of a protein. It predicts the damaging effects of any type of protein sequence variations to not only single amino acid substitutions but also in-frame insertions, deletions, and multiple amino acid substitutions. The results are obtained as either “Deleterious” if the prediction score was <-2.5, while score > −2.5 indicates that the variant is predicted to have a “Neutral” effect. (Available at: http://provean.jcvi.org/index.php).^(27, 28)^

#### 2.2.3. Polyphen-2 Server

Polymorphism Phenotyping v2.0 (PolyPhen-2) is another online tool that predicts the possible effects of an amino acid substitution on the structure and function of the protein. The results are classified into “PROBABLY DAMAGING” that is the most disease causing with a score near to 1 (0.7-1), “POSSIBLY DAMAGING” with a less disease causing ability with a score of 0.5-0.8 and “BENIGN” which does not alter protein functions with a score closer to zero; (0-0.4). (Available at: http://genetics.bwh.harvard.edu/pph2/).^(22, 29, 30)^

#### 2.2.4. SNPs & Go server

An online web server that used to ensure the disease relationship with the studied single nucleotide polymorphisms SNPs. It gives three different results based on three different analytical algorithms; Panther result, PHD-SNP result, SNPs&GO result. Each one of these results is composed of three parts, the prediction which decides whether the mutation is neutral or disease related, reliability index (RI), and disease probability (if >0.5 mutation is considered as disease causing nsSNP). (Available at: http://snps-and-go.biocomp.unibo.it/snps-and-go/).^(31)^

#### 2.2.5. PMUT Server

PMUT is a powerful web-based tool used for the prediction of pathological variants on proteins. The prediction results are classified as “Neutral” or “Disease”. It is available at (http://mmb.irbbarcelona.org/PMut).^(32)^

### 2.3. Protein Structural analysis

**I-Mutant:**

A online web-tool that is used for the prediction of protein stability changes upon single point mutations, determining whether the mutation increases or decreases the protein stability. (Available at: http://gpcr.biocomp.unibo.it/cgi/predictors/I-Mutant2.0/I-Mutant2.0.cgi).^(33)^

### 2.4. Project HOPE

It is an online web-server used to analyze the structural and functional variations on protein sequence that have been resulted from single amino acid substitution. It searches protein 3D structures by collecting structural information from a series of sources, including calculations on the 3D coordinates of the protein and sequence annotations from the UniProt database. Protein sequences are submitted to project HOPE server then HOPE builds a complete report with text, figures, and animations. It is available at (http://www.cmbi.ru.nl/hope).^(34)^

### 2.5. GeneMANIA

A user-friendly web interface tool approaches to know protein function, analyzing submitted gene lists and prioritizing genes for functional assays. It provides lists with functionally similar genes that it identifies and recognizes using available genomics and proteomics data. (Available at: (http://www.genemania.org/).^(35)^

### 2.6. Modeling nsSNP locations on protein structure

**Chimera:**

Chimera 1.8 software is used for visualization and editing of the three dimensional structure of protein. It visualizes the original amino acid with the mutated one to see the impact that can be produced. It is a highly extensible program for interactive visualization and analysis of molecular structures and related data, including density maps, supramolecular assemblies and sequence alignments. High-quality images and animations can be generated. (Availabe at: http://www.cgl.ucsf.edu/chimera).^(36)^

### 2.7. 3′-UTR analysis

#### 2.7.1. Ensemble 95

RNA sequence were retrieved from ensemble, which is website that makes key genomic data sets available to the entire scientific community without restrictions.^(37, 38)^ (Available at: https://www.ensembl.org/index.html).

#### 2.7.2. RegRNA2.0

Then the RNA sequences were inserted to RegRNA 2.0 (which is an integrated web server for identifying functional RNA motifs and sites.^(39, 40)^ (Available at: http://regrna2.mbc.nctu.edu.tw/) to find the related microRNA (miRNA).

#### 2.7.3. miRmap22.1

Then the miRNA sequence were further inserted in to miRmap website to predict the miRNA targets and study the repression strength using thermodynamic, probabilistic, evolutionary and sequence-based approaches.^(41)^ (Available at: https://mirmap.ezlab.org/).

#### 2.7.4. miRBase

After that we use mirbase website to study its annotations (miRBase website provides a wide-range of information on published microRNAs, including their sequences, their biogenesis precursors, genome coordinates, and community driven annotation).^(42–44)^ (Available at: http://www.mirbase.org/).

## Result

339 missense SNPs were retrived from National Center for Biotechnology Information (NCBI) website and submitted to SIFT, Proven, Polyphen-2, SNAP2 respectively. After analysis, 60 mutations out of 339 missense mutations were found to be deleterious in all four server. as found in table (1)

**Table. (1):** Shows the most deleterious SNPs by four softwares.

| dbSNP rs# | Substitution | sift Prediction | sift Score | Provean Prediction | Provean Score | POLYPHEN2 Prediction | POLYPHEN2 Score | SNAP2 Predicted Effect | SNA Scor |
| --- | --- | --- | --- | --- | --- | --- | --- | --- | --- |
| rs1229235381 | R5W | AFFECT | 0 | Deleterious | -2.538 | possibly damaging | 0.95 | Effect | 50 |
| rs1330567400 | C10R | AFFECT | 0 | Deleterious | -4.238 | probably damaging | 0.997 | Effect | 84 |
| rs1420986481 | L13P | AFFECT | 0 | Deleterious | -3.429 | probably damaging | 0.99 | Effect | 63 |
| rs867160323 | G19C | AFFECT | 0 | Deleterious | -4.108 | probably damaging | 1 | Effect | 85 |
| rs748586810 | N32T | AFFECT | 0 | Deleterious | -2.666 | probably damaging | 0.996 | Effect | 20 |
|  | N32I | AFFECT | 0 | Deleterious | -4.378 | probably damaging | 0.999 | Effect | 57 |
| rs1379715347 | V40E | AFFECT | 0.02 | Deleterious | -2.98 | possibly damaging | 0.828 | Effect | 35 |
| rs576105333 | Y61C | AFFECT | 0 | Deleterious | -2.61 | probably damaging | 1 | Effect | 71 |
| rs1224937342 | F68C | AFFECT | 0 | Deleterious | -2.561 | probably damaging | 0.992 | Effect | 31 |
| rs1356908272 | D69V | AFFECT | 0 | Deleterious | -3.745 | possibly damaging | 0.514 | effect | 76 |
| rs886055106 | R72W | AFFECT | 0 | Deleterious | -3.542 | probably damaging | 0.999 | effect | 85 |
| rs764744666 | Y86H | AFFECT | 0 | Deleterious | -2.834 | probably damaging | 0.999 | effect | 68 |
| rs889604735 | P89R | AFFECT | 0 | Deleterious | -3.725 | probably damaging | 1 | effect | 19 |
| rs1220794681 | E98G | AFFECT | 0 | Deleterious | -3.449 | probably damaging | 0.999 | effect | 50 |
| rs754648178 | W129R | AFFECT | 0 | Deleterious | -5.379 | probably damaging | 0.999 | effect | 69 |
| rs1302837695 | K167T | AFFECT | 0 | Deleterious | -3.663 | probably damaging | 0.994 | effect | 67 |
| rs377364475 | R178W | AFFECT | 0 | Deleterious | -3.983 | possibly damaging | 0.95 | effect | 87 |
| rs1157407921 | G181C | AFFECT | 0 | Deleterious | -2.715 | probably damaging | 1 | effect | 57 |
| rs750319630 | V186D | AFFECT | 0.01 | Deleterious | -3.175 | probably damaging | 0.969 | effect | 56 |
| rs1181193053 | A191E | AFFECT | 0 | Deleterious | -2.86 | possibly<br>damaging | 0.948 | effect | 56 |
| rs778114154 | P197L | AFFECT | 0 | Deleterious | -7.417 | probably<br>damaging | 1 | effect | 21 |
| rs778975572 | L204P | AFFECT | 0 | Deleterious | -4.144 | possibly<br>damaging | 0.772 | effect | 78 |
| rs769602895 | G222R | AFFECT | 0 | Deleterious | -4.367 | probably<br>damaging | 0.979 | effect | 84 |
| rs1422292058 | L224P | AFFECT | 0 | Deleterious | -4.894 | probably<br>damaging | 0.999 | effect | 77 |
| rs1474908539 | S230P | AFFECT | 0 | Deleterious | -2.73 | possibly<br>damaging | 0.718 | effect | 45 |
| rs748887234 | R234I | AFFECT | 0 | Deleterious | -4.45 | probably<br>damaging | 0.997 | effect | 68 |
| rs369174398 | Y236C | AFFECT | 0 | Deleterious | -5.723 | possibly<br>damaging | 0.927 | effect | 19 |
| rs1476963715 | G253E | AFFECT | 0 | Deleterious | -3.683 | probably<br>damaging | 1 | effect | 76 |
| rs1015369709 | C256R | AFFECT | 0 | Deleterious | -3.356 | possibly<br>damaging | 0.899 | effect | 39 |
|  | C256G | AFFECT | 0 | Deleterious | -3.232 | possibly<br>damaging | 0.718 | effect | 37 |
| rs778451530 | P269L | AFFECT | 0 | Deleterious | -5.08 | probably<br>damaging | 1 | effect | 68 |
| rs1439042141 | G271V | AFFECT | 0 | Deleterious | -4.416 | probably<br>damaging | 1 | effect | 60 |
| rs757839826 | P282S | AFFECT | 0 | Deleterious | -5.005 | probably<br>damaging | 1 | effect | 50 |
| rs1292011900 | P283L | AFFECT | 0 | Deleterious | -4.623 | probably<br>damaging | 0.959 | effect | 38 |
| rs746899310 | L285R | AFFECT | 0 | Deleterious | -4.499 | probably<br>damaging | 1 | effect | 42 |
| rs1463889939 | W302G | AFFECT | 0 | Deleterious | -10.032 | probably<br>damaging | 1 | effect | 87 |
| rs1412078281 | H318R | AFFECT | 0 | Deleterious | -3.987 | probably<br>damaging | 0.98 | effect | 30 |
| rs1319790255 | D324G | AFFECT | 0 | Deleterious | -4.179 | probably<br>damaging | 1 | effect | 3 |
| rs78488864 | W344R | AFFECT | 0 | Deleterious | -3.279 | probably<br>damaging | 0.999 | effect | 26 |
| rs1439891137 | W344C | AFFECT | 0 | Deleterious | -2.985 | probably<br>damaging | 1 | effect | 39 |
| rs752287637 | D350H | AFFECT | 0 | Deleterious | -5.89 | probably<br>damaging | 1 | effect | 83 |
| rs764182538 | H357Y | AFFECT | 0 | Deleterious | -2.596 | probably<br>damaging | 0.998 | effect | 25 |
| rs890180094 | H357P | AFFECT | 0 | Deleterious | -6.354 | probably<br>damaging | 1 | effect | 90 |
| rs764124723 | T402I | AFFECT | 0 | Deleterious | -4.114 | probably<br>damaging | 1 | effect | 49 |
| rs751732245 | R406W | AFFECT | 0 | Deleterious | -6.971 | probably<br>damaging | 1 | effect | 75 |
| rs1211965571 | D416H | AFFECT | 0 | Deleterious | -2.771 | probably<br>damaging | 1 | effect | 25 |
| rs1288356140 | D416G | AFFECT | 0 | Deleterious | -3.396 | probably<br>damaging | 1 | effect | 12 |
|  | D416V | AFFECT | 0 | Deleterious | -4.712 | probably<br>damaging | 1 | effect | 8 |
| rs139061039 | D418G | AFFECT | 0 | Deleterious | -5.105 | probably<br>damaging | 1 | effect | 32 |
| rs759392662 | Y429C | AFFECT | 0 | Deleterious | -7.379 | probably<br>damaging | 1 | effect | 40 |
| rs1227872471 | D431H | AFFECT | 0 | Deleterious | -4.105 | probably<br>damaging | 1 | effect | 54 |
| rs1256723119 | L433S | AFFECT | 0 | Deleterious | -4.322 | probably<br>damaging | 1 | effect | 42 |
| rs766858148 | G448R | AFFECT | 0 | Deleterious | -3.596 | probably<br>damaging | 1 | effect | 48 |
| rs1409820369 | D473H | AFFECT | 0 | Deleterious | -4.896 | probably<br>damaging | 1 | effect | 52 |
| rs1279189246 | H478R | AFFECT | 0 | Deleterious | -3.431 | probably<br>damaging | 1 | effect | 68 |
|  | H478L | AFFECT | 0 | Deleterious | -5.421 | probably<br>damaging | 1 | effect | 62 |
| rs1333368709 | D483G | AFFECT | 0 | Deleterious | -5.332 | probably<br>damaging | 0.995 | effect | 62 |
| rs753532618 | R484G | AFFECT | 0 | Deleterious | -3.653 | probably<br>damaging | 0.996 | effect | 76 |
| rs1387898294 | D485E | AFFECT | 0 | Deleterious | -2.773 | probably<br>damaging | 0.971 | effect | 3 |
| rs749237410 | F498V | AFFECT | 0 | Deleterious | -4.135 | probably<br>damaging | 1 | effect | 55 |

Furthermore, the above result were submitted to SNP&GO, PHD, and Pmut to study the relation between the SNPs and the disease. Out of 60 SNPs 10 were found to be related to disease. As indicated in table (2). Then I mutant was used to detect the stability of the protein and the result indicated that all the SNPs except one (rs1279189246,H478L) lead to decrease in the stability which lead to unstable amino acid interaction. As found in table (3).

**Table. (2):** Disease effect nsSNPs associated variations predicted by SNPs&GO and PHD-SNP softwares.

| dbSNP rs# | SUB | SNP & GO | RI* | Probability | PHD | RI* | Probability | PMUT Score | Prediction |
| --- | --- | --- | --- | --- | --- | --- | --- | --- | --- |
| rs1356908272 | D69V | Disease | 3 | 0.648 | Disease | 8 | 0.893 | 0.68 (85%) | Disease |
| rs886055106 | R72W | Disease | 3 | 0.625 | Disease | 7 | 0.868 | 0.53 (80%) | Disease |
| rs750319630 | V186D | Disease | 0 | 0.509 | Disease | 7 | 0.832 | 0.64 (84%) | Disease |
| rs778114154 | P197L | Disease | 3 | 0.649 | Disease | 8 | 0.921 | 0.84 (91%) | Disease |
| rs778975572 | L204P | Disease | 1 | 0.533 | Disease | 7 | 0.825 | 0.68 (85%) | Disease |
| rs1422292058 | L224P | Disease | 1 | 0.535 | Disease | 7 | 0.836 | 0.86 (91%) | Disease |
| rs1256723119 | L433S | Disease | 1 | 0.539 | Disease | 7 | 0.828 | 0.86 (91%) | Disease |
| rs1279189246 | H478R | Disease | 1 | 0.564 | Disease | 8 | 0.913 | 0.70 (86%) | Disease |
| rs1333368709 | D483G | Disease | 0 | 0.518 | Disease | 8 | 0.885 | 0.86 (91%) | Disease |
| rs749237410 | F498V | Disease | 2 | 0.62 | Disease | 8 | 0.886 | 0.69 (86%) | Disease |

\* RI: Reliability Index

**Table. (3):** Stability analysis predicted by I-Mutant version 3.0.

| dbSNP rs# | Mutation | i-mutant prediction | RI | Score |
| --- | --- | --- | --- | --- |
| rs1356908272 | D69V | Decrease | 5 | -0.6 |
| rs886055106 | R72W | Decrease | 5 | -0.86 |
| rs750319630 | V186D | Decrease | 7 | -1.22 |
| rs778114154 | P197L | Decrease | 8 | -0.84 |
| rs778975572 | L204P | Decrease | 7 | -1.72 |
| rs1422292058 | L224P | Decrease | 4 | -1.5 |
| rs1256723119 | L433S | Decrease | 9 | -2.29 |
| rs1279189246 | H478L | Increase | 5 | 0.4 |
| rs1333368709 | D483G | Decrease | 5 | -1.02 |
| rs749237410 | F498V | Decrease | 9 | -1.76 |

\*RI: Reliability Index

One significant microRNA was found and investigated as shown in table (4) and (5).

**Table. (4):**
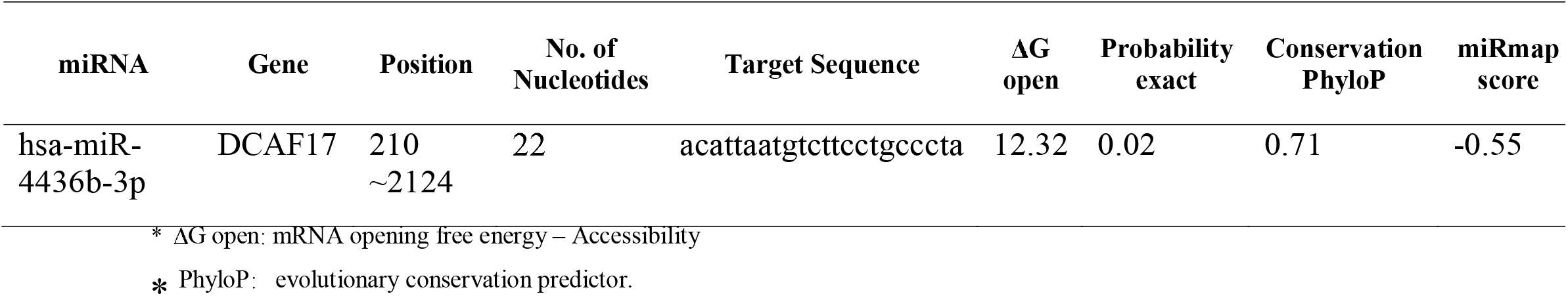
miRNAs target identified in 3′-UTR region of *DCAF17* gene by miRmap website.

**Table. (5):** Alignment of Query to mature miRNAs.

| Accession | ID | Query start | Query end | Subject start | Subject end | Strand | Score | Value |
| --- | --- | --- | --- | --- | --- | --- | --- | --- |
| MIMAT0019941 | hsa-miR-4436b-3p | 10 | 21 | 2 | 13 | - | 60 | 12 |
| MIMAT0025585 | mmu-miR-6540-5p | 2 | 22 | 2 | 22 | - | 60 | 12 |
| MIMAT0027651 | hsa-miR-6875-3p | 10 | 21 | 3 | 14 | + | 60 | 12 |
| MIMAT0017064 | mmu-miR-135a-2-3p | 4 | 22 | 3 | 21 | - | 59 | 15 |
| MIMAT0037309 | hsa-miR-135a-2-3p | 4 | 22 | 4 | 22 | - | 59 | 15 |
| MIMAT0015037 | hsa-miR-3163 | 9 | 21 | 10 | 22 | - | 56 | 27 |
| MIMAT0019772 | hsa-miR-4685-3p | 11 | 21 | 6 | 16 | + | 55 | 33 |
| MIMAT0027850 | mmu-miR-6974-5p | 6 | 19 | 5 | 18 | - | 52 | 58 |
| MIMAT0027973 | mmu-miR-7034-3p | 9 | 22 | 4 | 17 | + | 52 | 58 |
| MIMAT0028056 | mmu-miR-7075-5p | 7 | 20 | 2 | 15 | - | 52 | 58 |
| MIMAT0028057 | mmu-miR-7075-3p | 7 | 20 | 6 | 19 | + | 52 | 58 |
| MIMAT0019361 | hsa-miR-3976 | 10 | 21 | 6 | 17 | - | 51 | 70 |
| MIMAT0019940 | hsa-miR-4436b-5p | 10 | 21 | 8 | 19 | + | 51 | 70 |
| MIMAT0027746 | mmu-miR-6923-5p | 10 | 21 | 4 | 15 | - | 51 | 70 |
| MIMAT0003339 | hsa-miR-421 | 3 | 12 | 7 | 16 | - | 50 | 85 |
| MIMAT0004869 | mmu-miR-421-3p | 3 | 12 | 7 | 16 | - | 50 | 85 |

**Table. (6):** The gene co-expressed, share domain and interaction with *DCAF17* gene network.

| Gene 1 | Gene 2 | Weight | Network group |
| --- | --- | --- | --- |
| GEMIN2 | C18orf25 | 0.020875 | Co-expression |
| C18orf25 | DCAF17 | 0.01954 | Co-expression |
| POT1 | CUL4B | 0.015105 | Co-expression |
| POT1 | TMEM267 | 0.010743 | Co-expression |
| SKIV2L2 | DCAF17 | 0.004223 | Co-expression |
| VIPAS39 | BBS7 | 0.006134 | Co-expression |
| POLI | DCAF17 | 0.01365 | Co-expression |
| GEMIN2 | SKIV2L2 | 0.001957 | Co-expression |
| RAD17 | CUL4B | 0.009911 | Co-expression |
| RAD17 | POT1 | 0.005801 | Co-expression |
| RAD17 | VIPAS39 | 0.002685 | Co-expression |
| EYA3 | POT1 | 0.02177 | Co-expression |
| RAD17 | SKIV2L2 | 0.012174 | Co-expression |
| TMEM267 | DCAF17 | 0.008714 | Co-expression |
| POT1 | TMEM267 | 0.010902 | Co-expression |
| SKIV2L2 | DCAF17 | 0.010967 | Co-expression |
| SKIV2L2 | CUL4B | 0.014126 | Co-expression |
| SKIV2L2 | POT1 | 0.015277 | Co-expression |
| BBS7 | DCAF17 | 0.016686 | Co-expression |
| EXOC2 | BLMH | 0.015184 | Co-expression |
| GEMIN2 | CUL4B | 0.011145 | Co-expression |
| GEMIN2 | INTS7 | 0.011782 | Co-expression |
| RAD17 | SKIV2L2 | 0.013417 | Co-expression |
| CUL4B | DCAF17 | 0.009401 | Co-expression |
| TMEM267 | DCAF17 | 0.002878 | Co-expression |
| TMEM267 | CUL4B | 0.005961 | Co-expression |
| TMEM267 | C18orf25 | 0.003443 | Co-expression |
| SKIV2L2 | DCAF17 | 0.010745 | Co-expression |
| POLI | TMEM267 | 0.004744 | Co-expression |
| GEMIN2 | POT1 | 0.011051 | Co-expression |
| RAD17 | DCAF17 | 0.010133 | Co-expression |
| RAD17 | TMEM267 | 0.007081 | Co-expression |
| BLMH | SKIV2L2 | 0.011372 | Co-expression |
| C18orf25 | DCAF17 | 0.026408 | Co-expression |
| TMEM267 | DCAF17 | 0.016632 | Co-expression |
| SKIV2L2 | DCAF17 | 0.01629 | Co-expression |
| NAA15 | DCAF17 | 0.022654 | Co-expression |
| GEMIN2 | DCAF17 | 0.019081 | Co-expression |
| GEMIN2 | TMEM267 | 0.015272 | Co-expression |
| SKIV2L2 | POT1 | 0.003939 | Co-expression |
| BBS7 | DCAF17 | 0.021653 | Co-expression |
| GEMIN2 | TMEM267 | 0.0039 | Co-expression |
| GEMIN2 | POT1 | 0.006586 | Co-expression |
| TMEM267 | DCAF17 | 0.017157 | Co-expression |
| POT1 | DCAF17 | 0.018457 | Co-expression |
| POT1 | CUL4B | 0.016976 | Co-expression |
| INTS7 | PTTG1 | 0.009717 | Co-expression |
| POLI | DCAF17 | 0.024595 | Co-expression |
| POLI | POT1 | 0.015922 | Co-expression |
| GEMIN2 | POT1 | 0.014995 | Co-expression |
| POT1 | DCAF17 | 0.019193 | Co-expression |
| NAA15 | SKIV2L2 | 0.005278 | Co-expression |
| INTS7 | SKIV2L2 | 0.0093 | Co-expression |
| INTS7 | NAA15 | 0.011806 | Co-expression |
| GEMIN2 | CUL4B | 0.014478 | Co-expression |
| GEMIN2 | POT1 | 0.017858 | Co-expression |
| RAD17 | CUL4B | 0.018899 | Co-expression |
| RAD17 | GEMIN2 | 0.015564 | Co-expression |
| AGO1 | POT1 | 0.012863 | Co-expression |
| VIPAS39 | AGO1 | 0.006072 | Co-expression |
| EXOC2 | INTS7 | 0.01908 | Co-expression |
| RAD17 | POT1 | 0.027534 | Co-expression |
| RAD17 | VIPAS39 | 0.012628 | Co-expression |
| INTS7 | AGO1 | 0.019896 | Co-expression |
| GEMIN2 | PTTG1 | 0.002871 | Co-expression |
| RAD17 | CUL4B | 0.013971 | Co-expression |
| RAD17 | GEMIN2 | 0.012185 | Co-expression |
| INTS7 | SKIV2L2 | 0.006518 | Co-expression |
| BLMH | EYA3 | 0.014397 | Co-localization |
| POT1 | DCAF17 | 0.009539 | Co-localization |
| EYA3 | DCAF17 | 0.02755 | Co-localization |
| EYA3 | POT1 | 0.008808 | Co-localization |
| AGO1 | DCAF17 | 0.026643 | Co-localization |
| AGO1 | POT1 | 0.008327 | Co-localization |
| ARL15 | DCAF17 | 0.025506 | Co-localization |
| ARL15 | POT1 | 0.006414 | Co-localization |
| NAA15 | DCAF17 | 0.011778 | Co-localization |
| NAA15 | POT1 | 0.003766 | Co-localization |
| NAA15 | ARL15 | 0.008719 | Co-localization |
| INTS7 | DCAF17 | 0.023841 | Co-localization |
| INTS7 | POT1 | 0.008081 | Co-localization |
| INTS7 | NAA15 | 0.010854 | Co-localization |
| BLMH | DCAF17 | 0.025363 | Co-localization |
| BLMH | POT1 | 0.009025 | Co-localization |
| BLMH | EYA3 | 0.02518 | Co-localization |
| BLMH | NAA15 | 0.011865 | Co-localization |
| VIPAS39 | DCAF17 | 0.023599 | Co-localization |
| VIPAS39 | POT1 | 0.00637 | Co-localization |
| VIPAS39 | ARL15 | 0.025043 | Co-localization |
| VIPAS39 | NAA15 | 0.008351 | Co-localization |
| EXOC2 | DCAF17 | 0.024342 | Co-localization |
| EXOC2 | POT1 | 0.007643 | Co-localization |
| EXOC2 | ARL15 | 0.024647 | Co-localization |
| EXOC2 | VIPAS39 | 0.028187 | Co-localization |
| GEMIN2 | DCAF17 | 0.012483 | Co-localization |
| GEMIN2 | POT1 | 0.004406 | Co-localization |
| GEMIN2 | BLMH | 0.01327 | Co-localization |
| RAD17 | DCAF17 | 0.015909 | Co-localization |
| RAD17 | POT1 | 0.005233 | Co-localization |
| RAD17 | EYA3 | 0.015112 | Co-localization |
| PTTG1 | DCAF17 | 0.200927 | Genetic Interactions |
| NAA15 | SKIV2L2 | 0.060909 | Physical Interactions |
| CUL4B | DCAF17 | 0.115469 | Physical Interactions |
| RAB40B | DCAF17 | 0.353314 | Physical Interactions |
| ZMAT3 | DCAF17 | 0.300627 | Physical Interactions |
| BLMH | CUL4B | 0.065557 | Physical Interactions |
| POLI | CUL4B | 0.13731 | Predicted |

**Figure. (1):**
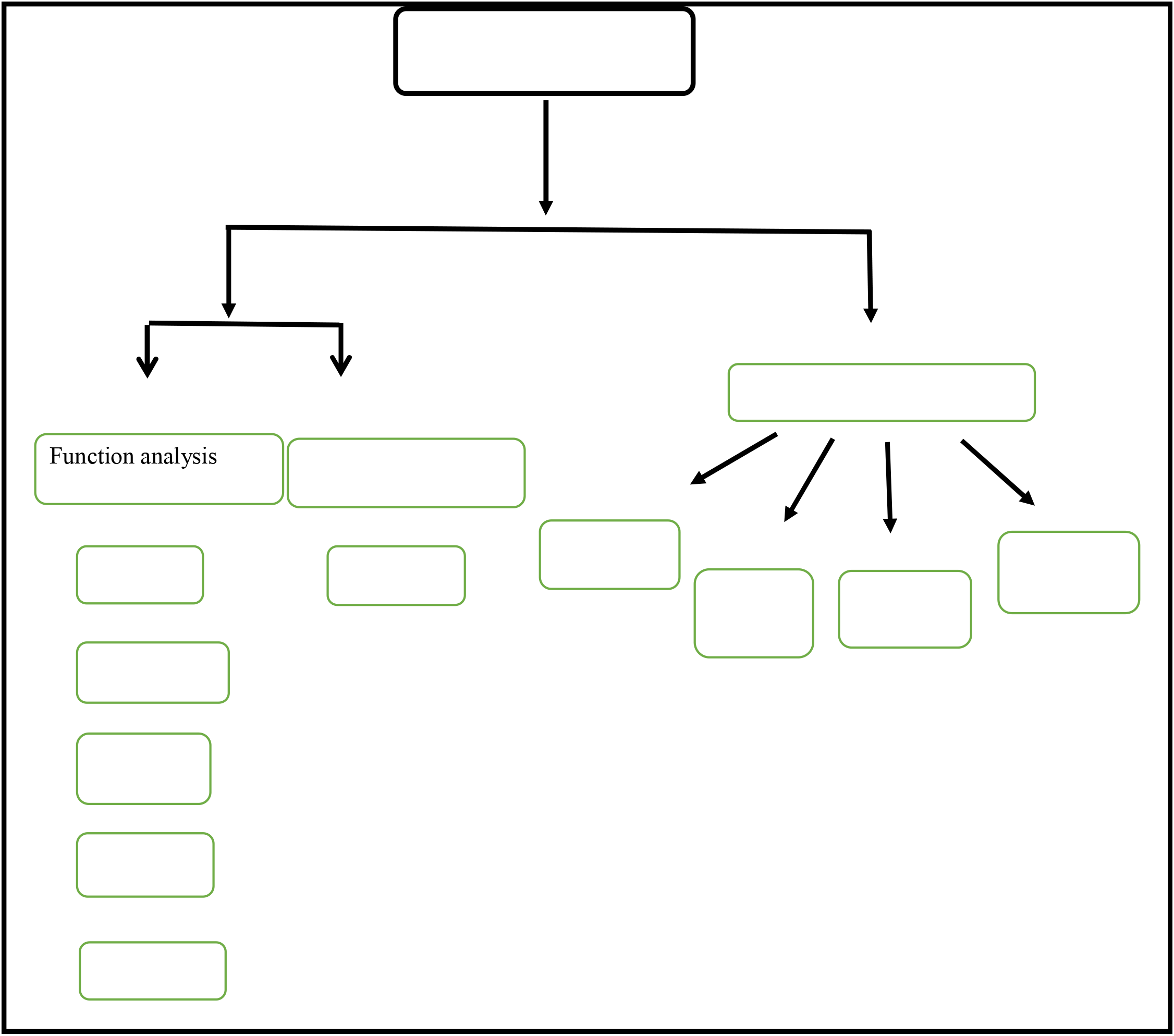
Diagrammatic representation of in silico work flow for *DCAF17* gene in coding and 3′-UTR regions.

**Figure. (2):**
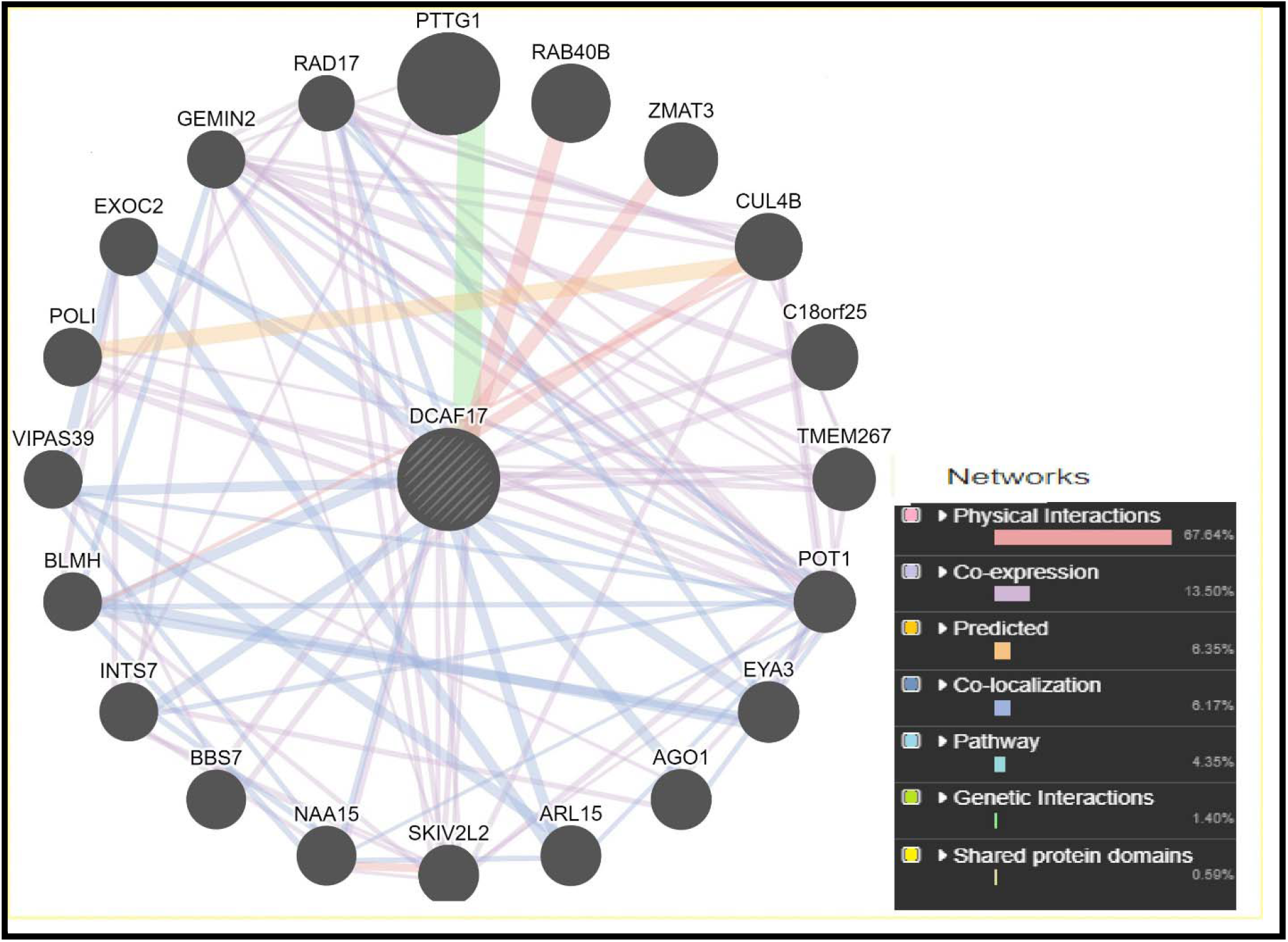
Interaction between *DCAF17* gene and related genes using GeneMANIA.

**Figure. (3):**
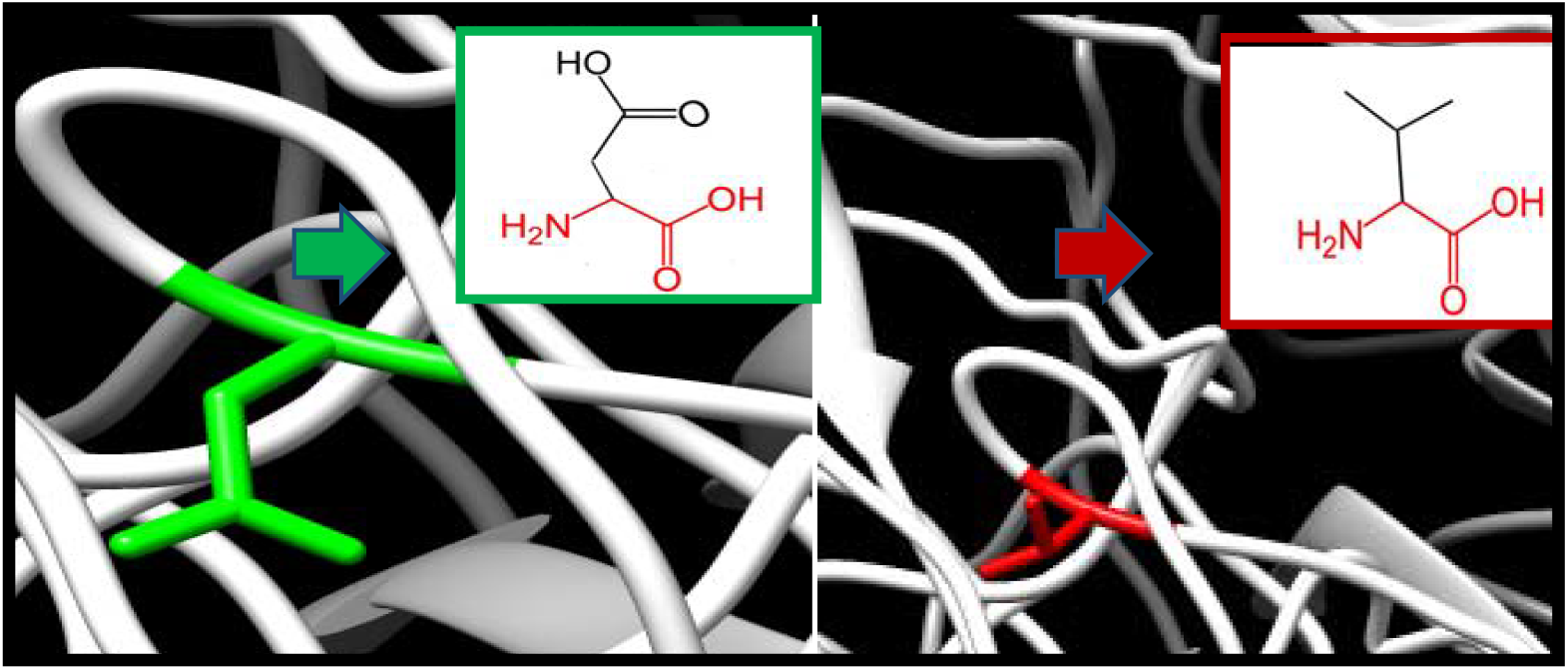
D69V: amino acid change from Aspartic acid to Valine at position 69.

**Figure. (4):**
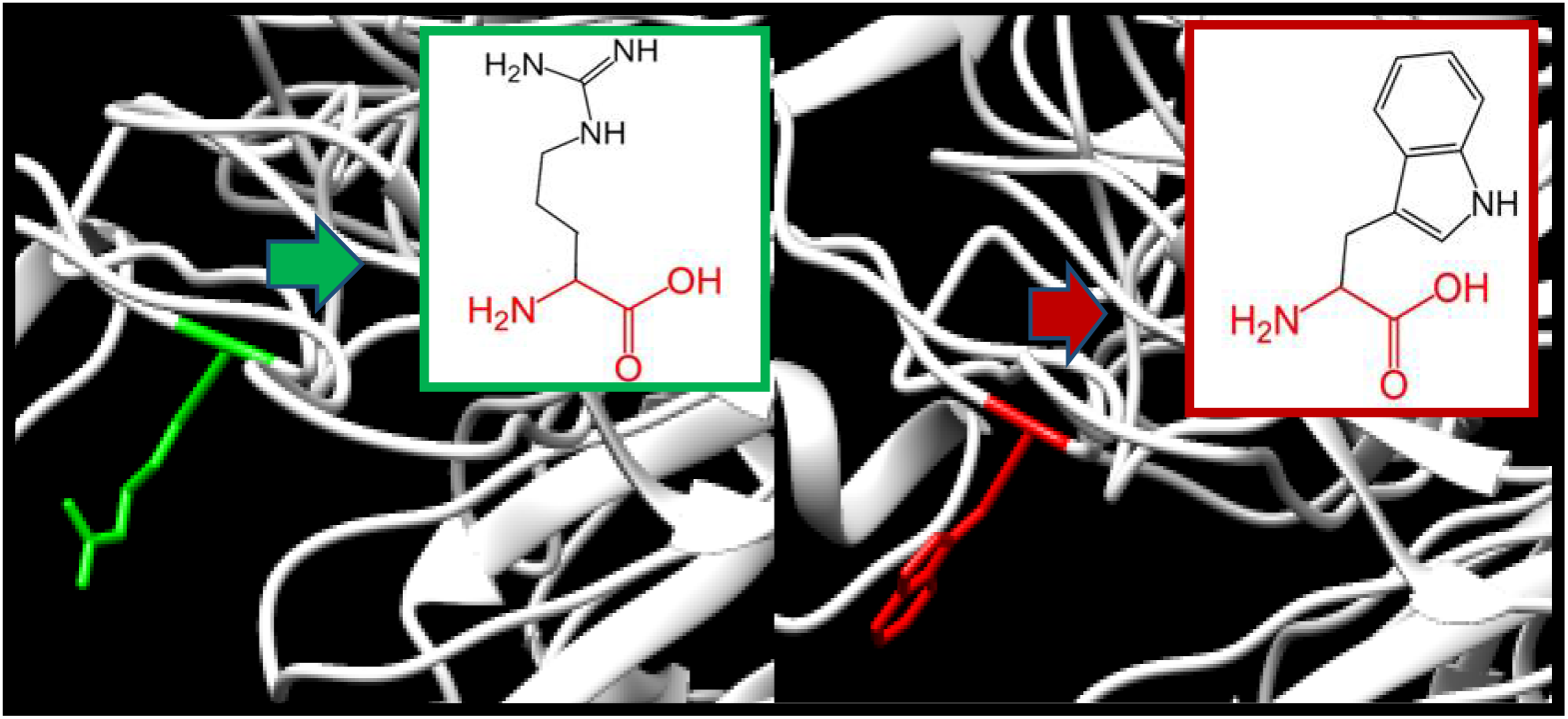
R72W: amino acid change from Arginine to Tryptophan at position 72.

**Figure. (5):**
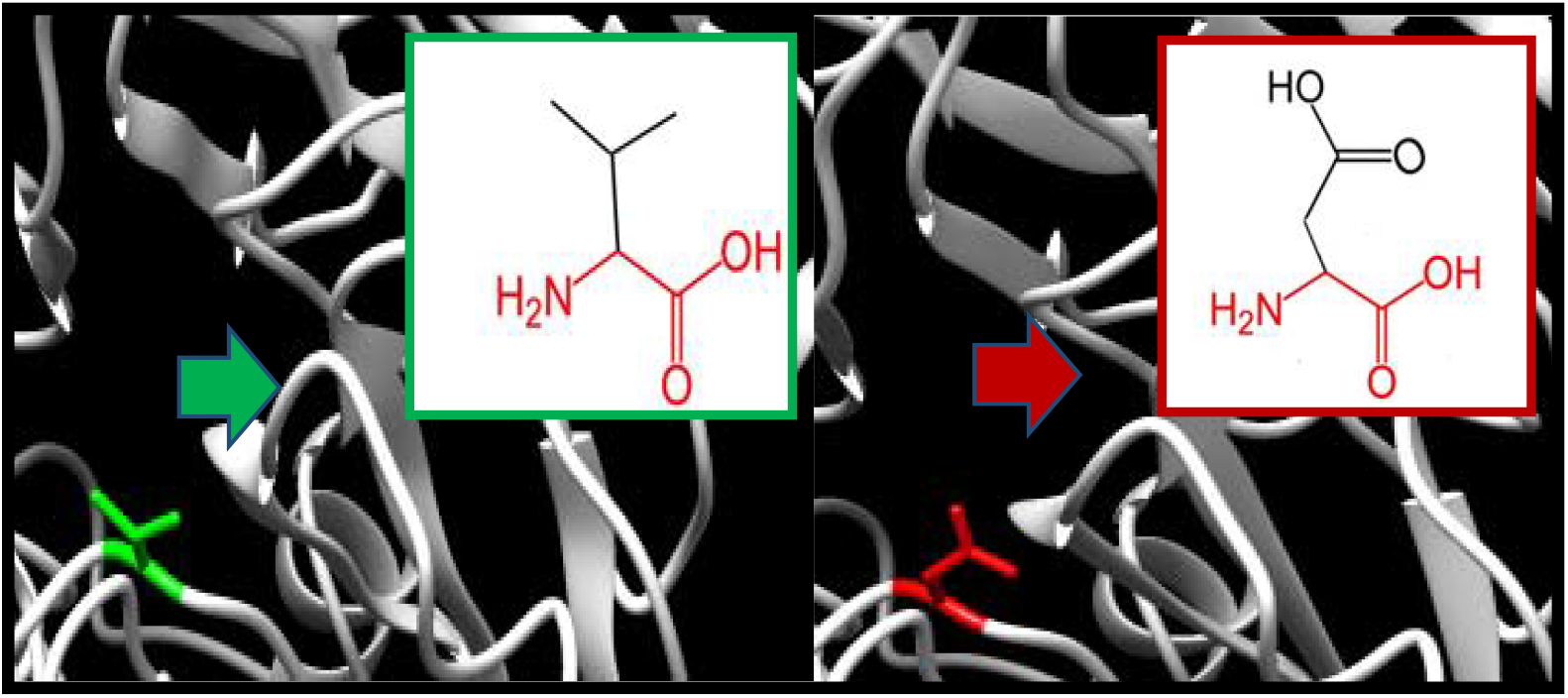
V186D: amino acid Valine change to Aspartic acid at position 186.

**Figure. (6):**
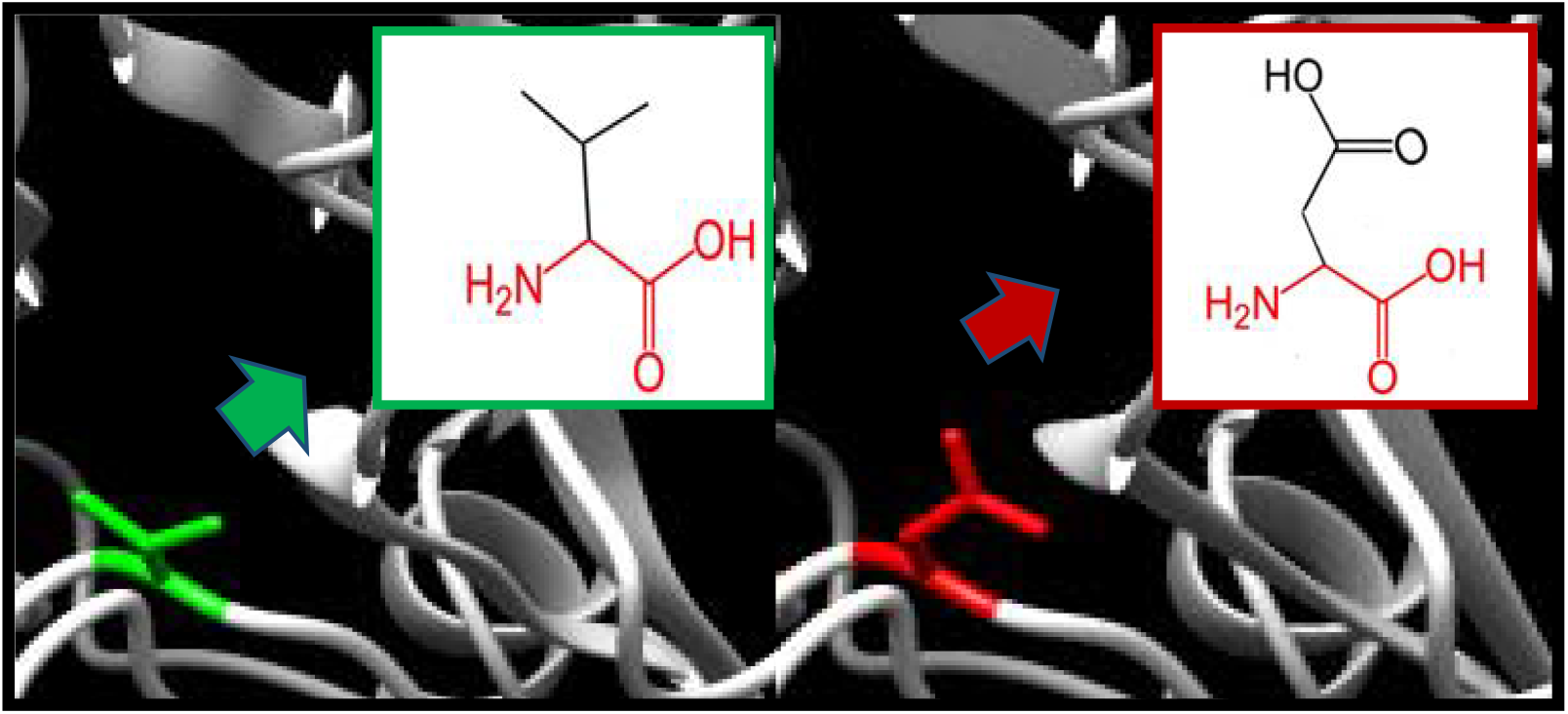
P197L: amino acid Proline change to Leucine at position 197.

**Figure. (7):**
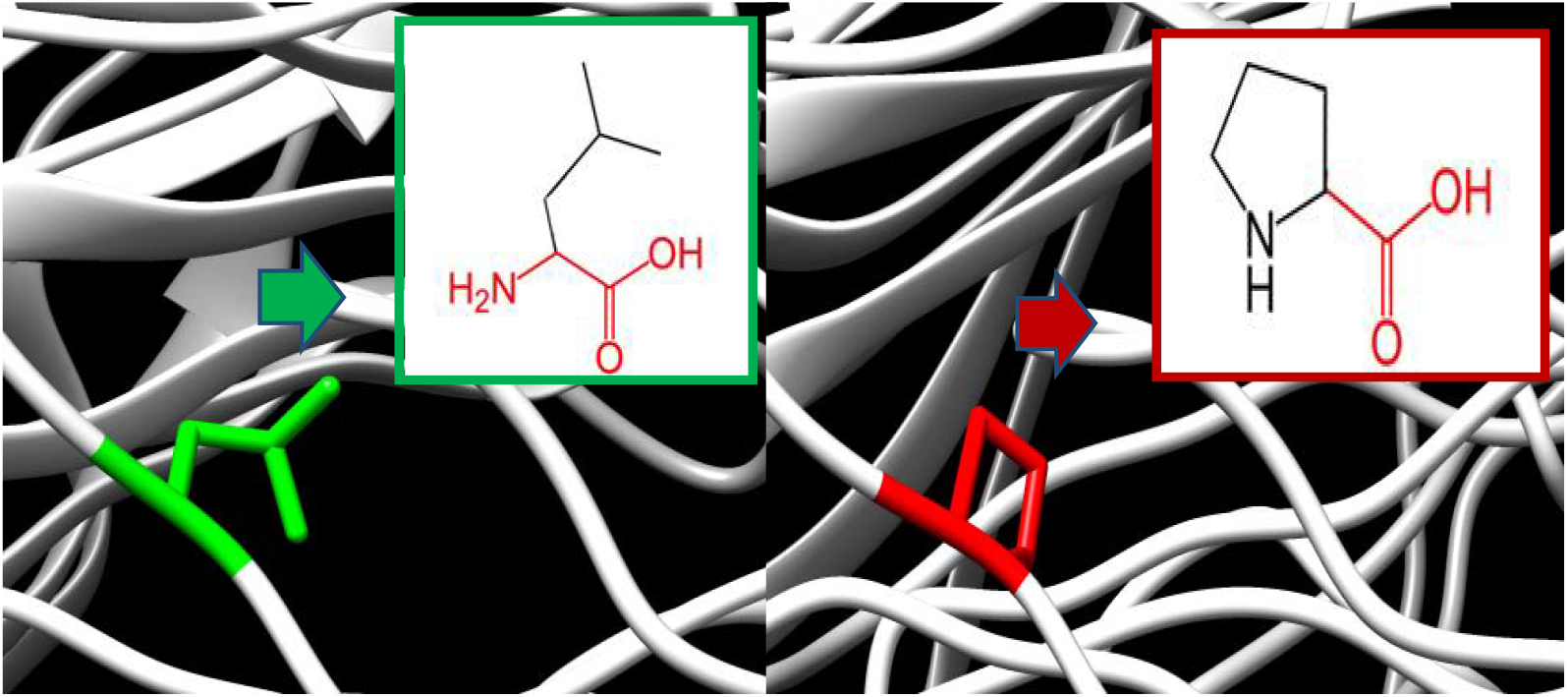
L204P: aminoacid Leucine change to Proline at position 204.

**Figure. (8):**
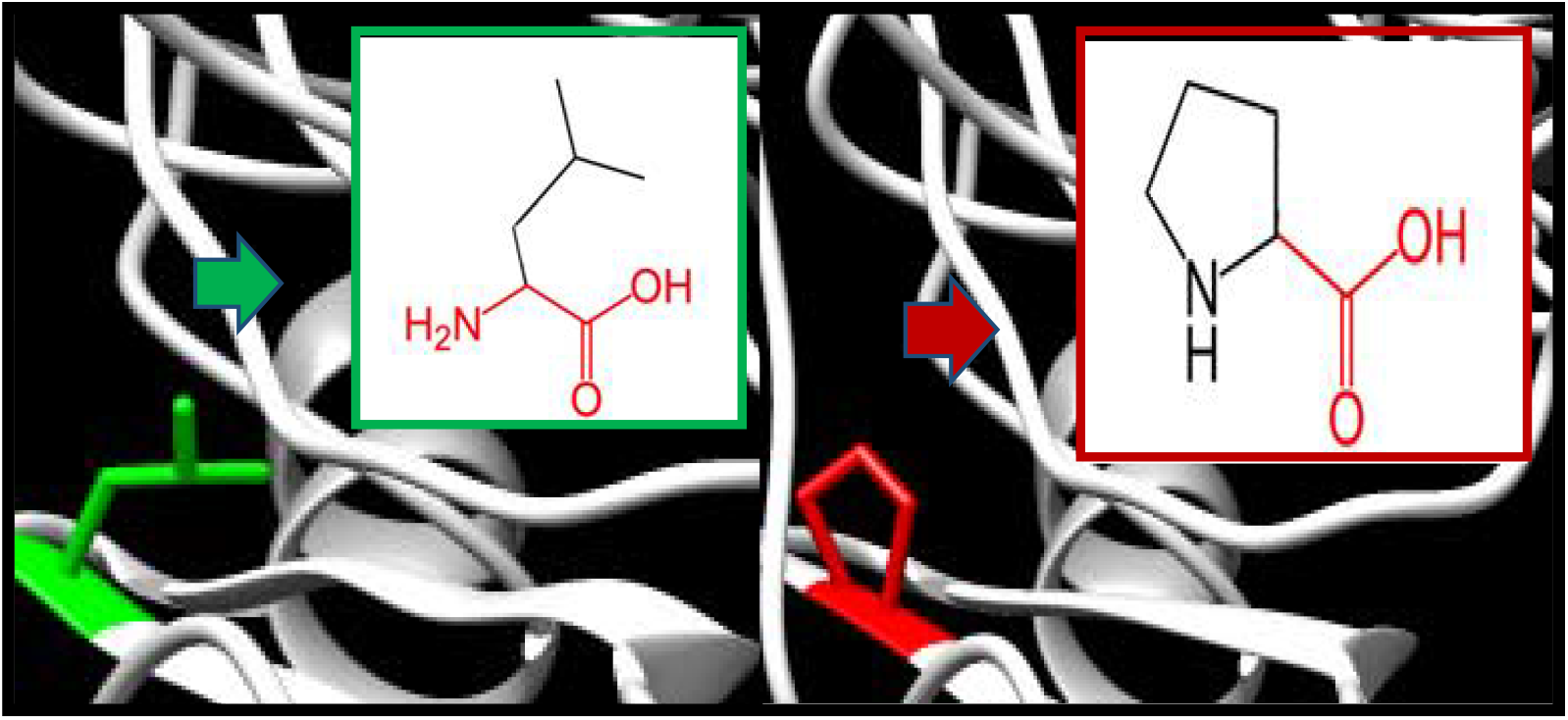
L224P: amino acid change from Leucine to Proline at position 224.

**Figure. (9):**
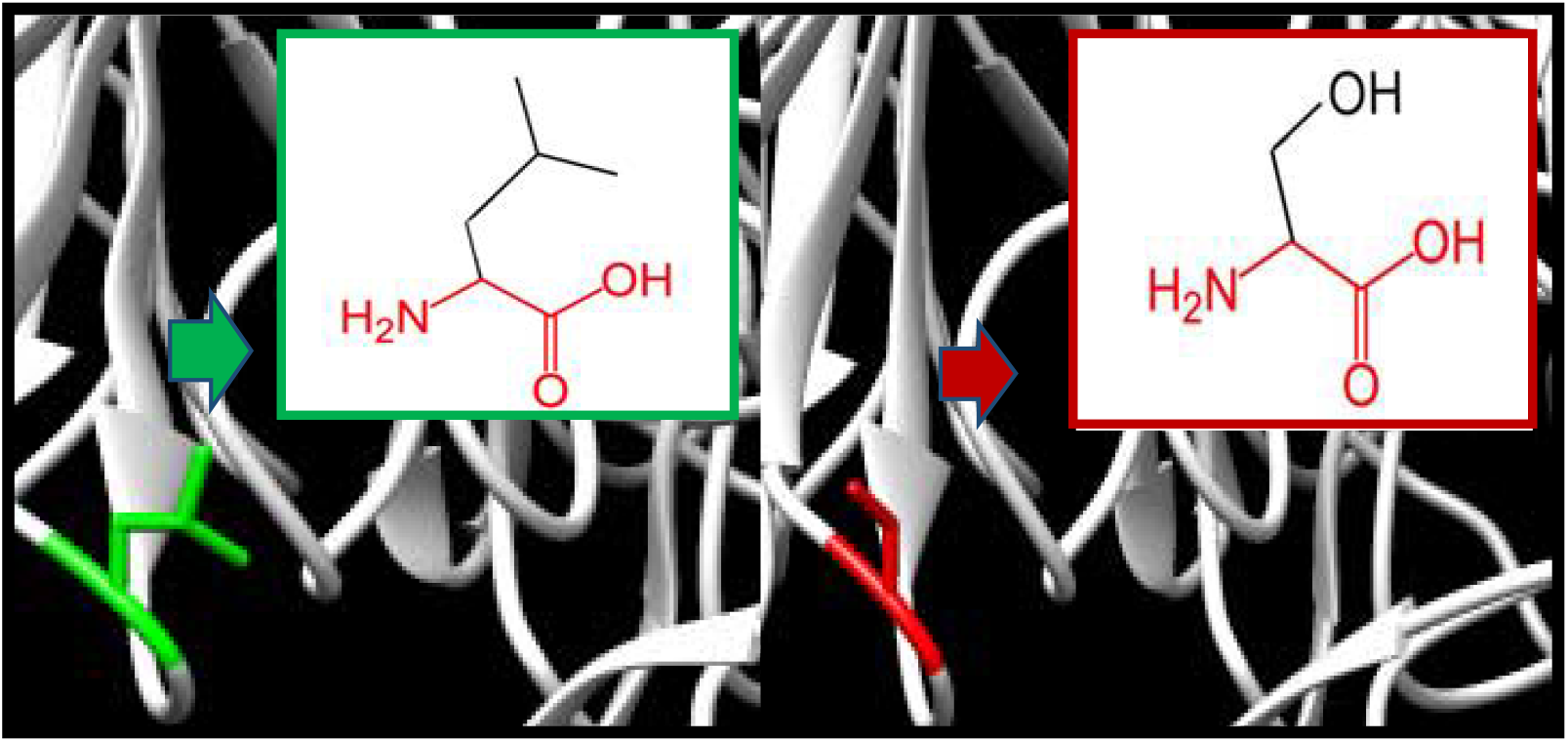
L433S: amino acid Leucine change to Serine at position 4333.

**Figure. (10):**
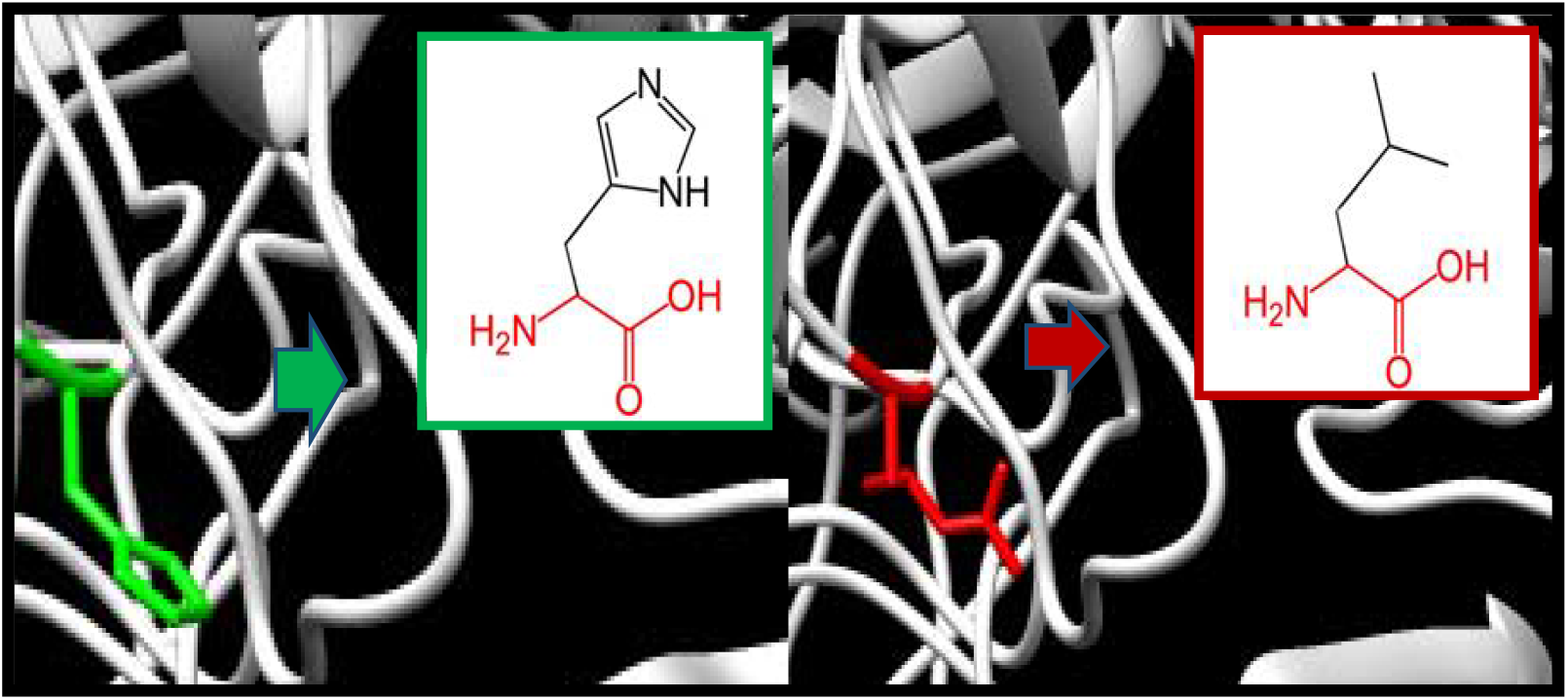
H478R: amino acid Histidine change to Arginine at position 478.

**Figure. (11):**
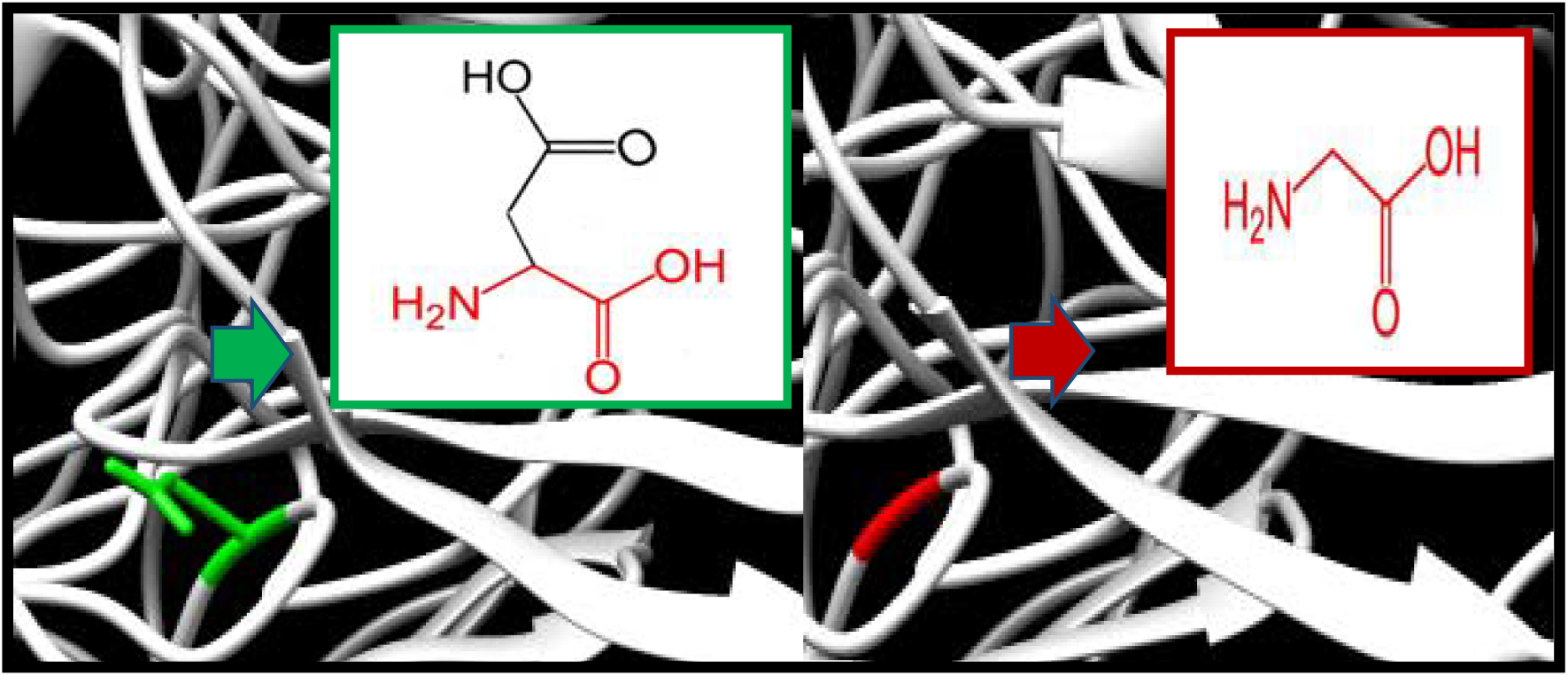
D483G: amino acid Aspartic acid change to Glycine at position 483.

**Figure. (12):**
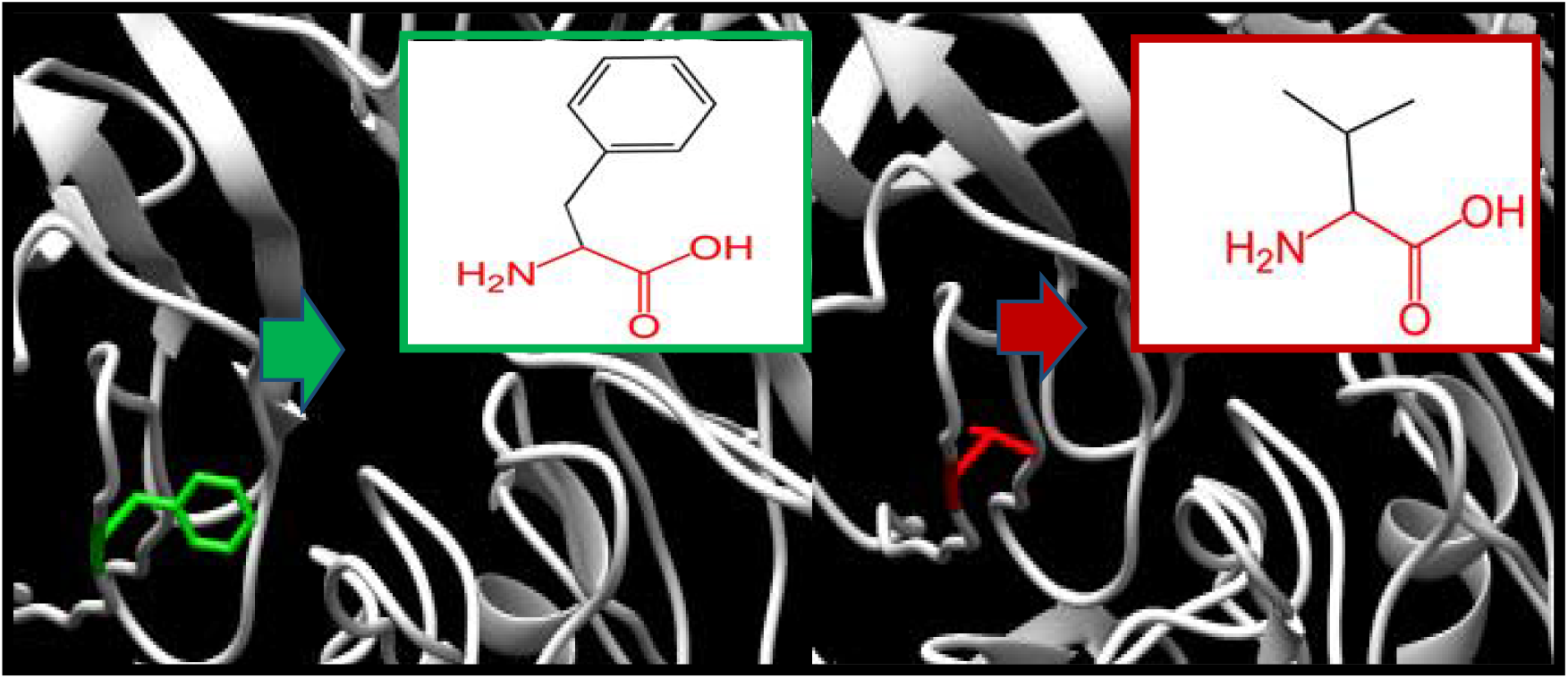
F498V: amino acid Phenylalanine change to Valine at position 498.

## Discussion

Ten novel mutations have been found that affect the stability, structure and the function of *DCAF17* gene with one associated microRNA in the 3′-UTR region utilizing various *in silico* bioinformatics tools. These methods for prediction used standard tools and multiple disciplines to detect and characterize the missense variation. In order to ensure our results and increase the accuracy; eight web server *in silico* algorithm have been used: SIFT, Proven, Polyphen, SNAP 2, SNP&GO, PHD, Pmut and I mutant.

This gene considered to be with no function but it has relation to the disease Woodhouse-Sakati Syndrome with MIM no 241080 (from NCBI)^(13, 45)^. Because there is no similar mutation pathogenicity prediction study to this gene; this founding considered to be novel mutation^(46), (13), (47)^.

GeneMANIA did not indicated specific function for *DCAF17* gene, but there is interaction with another genes^(47), (15), (48), (14)^. The function of the gene is not clear until now^(13)^.

Project HOPE server was used to predict the effect of the amino acid substitutions on the molecular level of the protein. The ten SNPs were submitted to hope server individually and the obtained result indicated a change in the size, hydrophobicity and charge of the final protein, all the SNPs were found in Ddb1- And Cul4-Associated Factor 17 IPR031620 domain region in the gene, which further confirm its pathogenicity.

All the mutant amino acids except three (R72W, V186D, P197L) were found to be smaller in size in comparison to the wild type which may lead to loss of interactions with other molecules or residues. The hydrophobicity of the wild-type and mutant residue differs, there were 6 SNPs mentioned in project hope report (D69V, R72W, V186D, L433P, H478L, D483G) that has mutant amino acid with more hydrophobicity that could lead to loss of hydrogen bonds and/or disturb correct folding. There is a difference in charge between the wild-type and mutant amino acid, with three SNPS(D69V, R72W, D483G) mentioned in the report to have different nerutral charge and one SNPs (V186D) with negative charge in the mutant amino acids which could cause a loss of interactions with other molecules or residues.

Different biological processes are controlled by transcriptional regulation done by non-coding RNA molecules (miRNA) consisting of 18–24 nucleotides in length which could activate and/or suppress protein translation inside the cells at post-transcriptional level.^(49–52)^ it was found in gene card site (https://www.genecards.org/) that *DCAF17* gene may be dysregulated by miRNA, so we analyzed the microRNA in the 3′-UTR region of *DCAF17* gene using multiple softwares (ensemble, regRNA2, miRmap and mirbase). We found one relevant miRNA; (hsa-miR-4436b-3p) with PhyloP conservation score of 0.71 and after analyzing its annotation through miRmap using sequence from 5 species human,mouse,worm,fly and Arabidopsis it appeared to have a role in inhibiting adipose triglyceride lipase, carcinogenesis and other possible roles.

This study was limited as it depended in computerized software and it need further wet lab studies for confirmation.

We now appreciate the regulatory role of the 3′-UTR in the function of DCAF17 protein and with this 10 novel mutation we could develop a precise biomarker for Woodhouse-Sakati Syndrome in the future.

## Conclusion

Ten novel SNPs were found to be deleterious in this paper with possible role as future biomarker for Woodhouse-Sakati syndrome, and one microRNA was found in the 3′-UTR to regulate the function of the protein.

## Acknowledgment

We appreciate the kind help and assistance we had in Africa City of Technology.

